# Characterizing massa intermedia morphology in schizophrenia: associations with aging, neuropsychological functioning, and atypical hippocampal development

**DOI:** 10.1101/2025.02.26.640398

**Authors:** Zachary Bergson, Maxwell J. Roeske, Baxter P. Rogers, Anna S. Huang, Victoria Fox, Stephan Heckers, Neil D. Woodward

## Abstract

The massa intermedia (MI) is a midline structure connecting the left and right thalamus that typically develops during the 2^nd^ trimester of pregnancy. Missing and smaller MI has been linked to neurodevelopmental disorders, including schizophrenia (SZ), and subtle deficits in cognition. However, findings are inconsistent and the association between MI and other anatomical variants linked to atypical brain development in SZ, including incomplete hippocampal inversion (IHI), is unclear. Presence/absence and morphology of the MI were ascertained on structural T1-weighted MRI images obtained at 3T in SZ (*n* = 223) and healthy individuals (*n* = 194) and compared between groups. Associations between MI morphology, cognitive function, and incomplete hippocampal inversion (IHI) were assessed. Prevalence of missing MI was 1.7% and did not differ between groups. MI was significantly smaller in SZ (*p* <.001). However, follow-up analyses revealed that smaller MI size in SZ was due to a significant Diagnosis x Age interaction characterized by a stronger negative age effect in SZ. IHI was significantly more common in individuals with missing MI. Neurocognition was not correlated with MI size when controlling for age and diagnosis. Stronger effects of age on MI size in SZ suggests that abnormal MI size measured in adulthood may not be a reliable static indicator of atypical neurodevelopment, but may reflect disease progression or accelerated aging. Missing MI was rare in our sample. Conversely, missing MI is associated with IHI suggesting a shared neurodevelopmental disruption in the 2^nd^ trimester.

## 1. Introduction

The massa intermedia (MI), also known as the interthalamic adhesion, is a midline structure connecting the left and right thalamus across the third ventricle that usually forms between 13-14 weeks of gestation (Rosales et al., 1968). There are considerable individual differences in the size of the MI, which ranges between 11.81mm-17.41mm and 8.47mm-12.47mm in the axial and coronal planes, respectively (Borghei et al., 2020). Moreover, approximately 4-25% of individuals in the general population are missing a MI (Allen and Gorski, 1991; Borghei et al., 2021; Damle et al., 2017; Nopoulos et al., 2001; Takahashi et al., 2017). The functional significance of the MI is unclear; however, it is well developed in lower mammals (Malobabić et al., 1990; Patra et al., 2022; Snyder et al., 1998) and is involved in interhemispheric transfer in rats and cats (Hirayasu and Wada, 1992; Hiyoshi and Wada, 1988a, 1988b). Neurodevelopmental disorders, including 6q termination deletion syndrome, Chiari II malformation, Cornelia de Lange syndrome, X-linked hydrocephalus, and pediatric midline brain abnormalities (Elia et al., 2006; Naidich et al., 1980; Whitehead et al., 2015; Whitehead & Najim, 2020; Yamasaki et al., 1995), are associated with abnormal MI morphology. Missing MI and smaller MI has also been linked to neuropsychological impairment in clinical populations and individual differences in cognition in healthy people (Borghei et al., 2020; Damle et al., 2017; Trzesniak et al., 2016; Vidal et al., 2024). Combined, these findings suggest missing MI and variability in its size may reflect atypical prenatal brain development.

Given the timing of its development, association with neurodevelopmental disorders, and potential linkage to cognitive function, the MI has attracted considerable attention among schizophrenia researchers as atypical prenatal brain development is one of the leading etiological theories of the disorder (Murray and Lewis, 1987; Weinberger, 1987). Consistent with the neurodevelopmental hypotheses, schizophrenia is associated with both smaller MI and higher prevalence of missing MI (Landin-Romero et al., 2016; Shimizu et al., 2008; Snyder et al., 1998; Takahashi, Suzuki, et al., 2008; Takahashi et al., 2017; Trzesniak et al., 2011, 2012). However, findings are mixed, especially with respect to missing MI. The inconsistent findings are likely due, at least in part, to small sample sizes and methodological differences across studies. The only meta-analysis investigating missing MI in schizophrenia (Trzesniak, et al., 2011), which found it to be higher in the disorder compared to healthy individuals (odds ratio = 1.98), included studies that used variable magnetic field strengths (1-3T), image resolutions, sample sizes (55-226 participants) and methods for measuring MI (Ceyhan et al., 2008; Erbaǧcı et al., 2002; Ettinger et al., 2007; Nopoulos et al., 2001; Shimizu et al., 2008; Snyder et al., 1998; Takahashi, Suzuki, et al., 2008). This resulted in a broad range of missing MI estimates in schizophrenia (4.7%-34.6%). Since this meta-analysis was published, only one of three subsequent studies replicated higher prevalence of missing MI (Landin-Romero et al., 2016; Takahashi et al., 2017; Trzesniak et al., 2012). Two of these studies examined MI length in schizophrenia (Takahashi et al., 2017; Trzesniak et al., 2012) and both found it to be smaller in the disorder compared to healthy individuals. Moreover, recent research showing a robust negative association between age and MI size in both the general population (Borghei, et al., 2021) and schizophrenia (Takahashi et al., 2017) raise concerns as to whether MI size measured in adulthood is a reliable marker of atypical brain development.

In this study, we used our large cohort of individuals with schizophrenia and healthy individuals that have undergone structural neuroimaging to: 1) characterize MI abnormalities in schizophrenia, including prevalence of missing MI and MI size; 2) if present, determine whether abnormal MI morphology measured in adulthood is a reliable indicator of disrupted early brain development (i.e. not a consequence of differential age effects); and 3) examine associations between MI characteristics and neuropsychological functioning. We also explored whether MI missingness and size are related to another indicator of brain development, incomplete hippocampal inversion (IHI), which also occurs during the second trimester and has been linked to schizophrenia (Roeske et al., 2021, 2022). An association between MI and IHI may provide additional evidence of disrupted brain development during the second trimester in schizophrenia. Research in children with and without midline brain malformations has shown a relationship between the two, as the presence of a hippocampal abnormality (IHI made up 96.3% of aberrations) more than tripled the odds of missing MI in one study (Whitehead and Najim, 2020).

## 2. Methods

### 2.1 Participants

This study included 223 individuals with schizophrenia spectrum disorders (SSD; 96 schizophreniform disorder, 86 schizophrenia, 41 schizoaffective disorder) and 194 healthy individuals (Table 1). Of note, the current sample overlaps with the sample reported in our previous study on IHI in schizophrenia (Roeske et al., 2021). SSD individuals and healthy people were drawn from a repository study, the Vanderbilt Psychiatric Genotype/Phenotype (PGPP) study, which is comprised of people that participated in one of three neuroimaging projects (CT00762866, R01MH070560, R01MH102266) conducted at Vanderbilt University Medical Center (VUMC). SSD individuals were recruited from inpatient and outpatient units at VUMC and healthy individuals were recruited from the Nashville and surrounding community via advertisement. The VUMC institutional review board approved the study. All participants were compensated for their time and provided written informed consent. Diagnoses were confirmed, or ruled out in the case of healthy individuals, using the Structured Clinical Interview for DSM-IV (First et al., 2002). Exclusion criteria included the following: significant medical or neurological illness, head injury, pregnancy, age under 15 or above 65, history of intellectual disability, substance abuse or dependence (at least 1 month in patients, lifetime in healthy individuals), and any MRI contraindications. Healthy individuals were excluded if they had any current or past psychiatric illnesses or a first-degree relative with a psychotic illness.

**Table 1.** Demographic and clinical characteristics (*N* = 417)^a^.

| | Schizophrenia Spectrum Disorder ($n = 223$) | Healthy Individuals ($n = 194$) |
| --- | --- | --- |
| *Age (years)* | 28.2 (11.3; 15-60) | 28.0 (10.3; 17-57) |
| *Race<sup>b</sup>* |  |  |
| Black ($n = 121$) | 75 (33.6%) | 46 (23.7%) |
| White ($n = 280$) | 140 (62.7%) | 140 (72.1%) |
| Asian ($n = 6$) | 1 (0.44%) | 5 (2.5%) |
| Other ($n = 8$) | 5 (2.2%) | 3 (1.5%) |
| American Indian ($n = 2$) | 2 (0.8%) | 0 |
| *Sex* |  |  |
| Male ($n = 279$) | 156 (69.9%) | 123 (63.4%) |
| Female ($n = 138$) | 67 (30.1%) | 71 (36.6%) |
| *Years of Education ($n = 391$)<sup>c</sup>* | 13.2 (2.26) | 14.9 (2.13) |
| *Chlorpromazine Equivalents<sup>d</sup> ($n = 198$)* | 407.6 (255.9) | -- |
| *WTAR<sup>†</sup> Standardized Score ($n = 398$)<sup>e</sup>* | 99.4 (16.2) | 111 (11.13) |
| *Incomplete Hippocampal Inversion<sup>f</sup>* | 64 (28.7%) | 33 (17.0%) |
| Left IHI score<sup>g</sup> | 2.92 (1.65) | 2.56 (1.19) |
| Right IHI score | 2.17 (1.10) | 1.98 (0.87) |
| *SCIP<sup>†</sup> Global Cognition Score ($n = 410$)<sup>h</sup>* | -1.14 (0.95) | 0.05 (0.65) |

<sup>a</sup> Table provides mean (standard deviation), (standard deviation; range), or count (%)
<sup>b</sup> Chi-square test of independence identified a significant difference between groups ( $p = .038$ )
<sup>c</sup> Mann-Whitney U test identified a significant difference between groups ( $p < .001$ )
<sup>d</sup> unit = mg
<sup>e</sup> Mann-Whitney U test identified a significant difference between groups ( $p < .001$ )
<sup>f</sup> Chi-square test of independence identified a significant difference between groups ( $p = .005$ )
<sup>g</sup> Mann-Whitney U test identified a significant difference between groups ( $p = .012$ )
<sup>h</sup> Mann-Whitney U test identified a significant difference between groups ( $p < .001$ )
<sup>†</sup> WTAR: Wechsler Test of Adult Reading; SCIP: Screen for Cognitive Impairment in Psychiatry (z-score)

### 2.2 MRI acquisition and processing

Structural MRI images were obtained at 3T on a Philips Intera Acheiva at Vanderbilt University Institute of Imaging Sciences. Each participant received a 3D T1-weighted scan (voxel resolution: 1 mm^3^; field of view = 256^2^; number of slices = 170; TE = 3.7 ms; TR = 8.0 ms). Images were visually inspected for excessive motion or artifacts before included in the analysis.

### 2.3 Assessment of massa intermedia presence and size

Massa intermedia presence and size were determined by identifying axial and coronal slices containing the third ventricle and using protocols described previously (Borghei et al., 2020; Borghei, et al., 2021; Vidal et al., 2024). ITK-SNAP version 4.0.1 (Yushkevich et al., 2006) was used for MI identification and measurement. Presence/absence and size of MI in the axial and coronal planes were assessed, blind to diagnosis, by two raters (ZB and NDW) after training. ZB evaluated MI morphology for all 417 participants. Presence of tissue connecting the left and right thalami were then examined in each slice depicting the midpoint of the third ventricle. If bridging tissue was identified in at least one slice in both axial and coronal views, the MI was considered present. A video describing MI identification is available to the public (https://doi.org/10.13140/RG.2.2.35022.47689). If the MI was deemed present, axial and coronal plane dimensions were measured at the midpoint of the third ventricle by identifying the longest slice of the MI in axial and coronal views. 25 participants were randomly selected to evaluate interrater reliability between ZB and NW. There was 100% agreement on MI presence between the two raters. Intraclass correlation coefficients (ICC) were estimated for MI measurements using a two-way random effects model based on single-item measurement. The ICCs for axial and coronal plane measurements were excellent (ICC = .932) and good (ICC = .892), indicating a high level of reliability between raters.

### 2.4 Neuropsychological Assessment

Neuropsychological functioning was assessed with the Screen for Cognitive Impairment in Psychiatry (SCIP; (Purdon, 2005)). The SCIP includes five subtests: verbal memory (immediate and delayed), working memory, verbal fluency, and processing speed. The SCIP is a reliable screening tool for quantifying cognition in psychotic disorders (Rojo et al., 2010) and has been validated in structural neuroimaging studies (Moussa-Tooks et al., 2024; Woodward and Heckers, 2015). Raw scores for each subscale were converted to z-scores using normative data and a composite SCIP z-score was then calculated by averaging subscale z-scores. The Wechsler Test of Adult Reading (standard score; (Wechsler, 2001)) was used to assess premorbid IQ.

### 2.5 Assessment of IHIs

Assessment of the presence and severity of IHI in this cohort has been described previously (see Roeske et al. 2021). The method for determining IHI was established in a study by Cury et al. (2015) in 2089 participants. Each hippocampus was given an IHI score ranging from 0 to 10. Scores of ≥3.75 met criteria for IHI. ITK-SNAP (version 3.8.0) was used to obtain IHI scores in the coronal view.

### 2.6 Statistical analyses

Statistical analyses were conducted in R version 4.3.3 (R Core Team, 2021) and jamovi version 2.3.28.0 (Şahin and Aybek, 2019). Chi-square tests of independence, Mann-Whitney U tests, and one-way analysis of variance were used to determine differences between groups in demographic characteristics. Prevalence of missing MI was compared between groups using a Chi-square test. Group differences in length of the MI in the axial and coronal planes were tested separately using step-wise linear regression. Step 1 included age, sex, and diagnosis as main effects. Given robust evidence of age effects on MI size and evidence of more pronounced age effects on brain structure/networks in SSD (Borghei et al., 2021; Cropley et al., 2017; Schnack et al., 2016; Sheffield et al., 2019; Takahashi et al., 2017), Step 2 added a Diagnosis x Age interaction. Model fit was examined by comparing change in adjusted R^2^. 5,000 sample bootstrap confidence intervals were generated to account for the negative skew in MI size when examining post-hoc comparisons between groups. Confidence intervals that did not contain zero were considered significant.

Linear regressions were also run in step-wise fashion to examine the effect of MI size on neurocognition. Step 1 included age, MI size, and diagnosis. This was followed by Step 2, which included a Diagnosis x MI size interaction. Model fit was examined by comparing change in adjusted R^2^. The SCIP subscales were examined in separate models if a significant effect of MI size was found for global cognition.

Associations between IHI and MI were tested several ways. First, a chi-square analysis was performed to determine if the frequency of IHI was associated with missing MI. Second, an analysis was performed to determine if the presence of IHI was associated with MI size. This was accomplished by adding a third step (Step 3: MI size ∼ Sex + Age + Diagnosis + Diagnosis x Age + IHI) to the step-wise linear regression described in the first paragraph of this section, with IHI presence added as a main effect. Finally, spearman correlations were used to explore associations between IHI severity scores (left and right) and MI size.

## 3. Results

### 3.1 Massa intermedia missingness and size

The prevalence of missing MI was low in our sample (1.7%, 7 participants) and did not differ between groups (Table 2). The initial step-wise linear regression (Step 1, Table 3), which included age, sex, and diagnosis as independent variables and MI lengths as dependent variables, revealed significant effects of age (*p* < .001, *d* = -0.81) and diagnosis (*p* < .001, *d* = -0.49) for axial length. Older age and SSD diagnosis were associated with smaller axial length. Sex was not a significant effect in the axial plane (*p* = .071, *d* = -0.18). Similar results were observed for coronal length (age: *p* < .001, *d* = -0.64 ; diagnosis: *p* < .001, *d =* -0.35); however, the sex effect was stronger and significant in the coronal plane (*p* = .001, *d* = -0.31), with male sex associated with smaller MI length.

**Table 2.** Massa Intermedia Size and Missingness (*n* = 415)^a^.

| | SZ ($n = 223$) | HC ($n = 192$) | $p$ | $r_{rb}$ |
| --- | --- | --- | --- | --- |
| Axial Length (mm) | 13.8 (3.77; 1.83-19.2) | 15.42 (2.94; 3.89-22.6) | <.001 | .252 |
| Coronal Length (mm) | 9.52 (2.55; 1.0-15.6) | 10.39 (2.33; 3.48-16.9) | <.001 | .208 |
| Missing Massa Intermedia | 4 (1.8%) | 3 (1.6%) | .855 | -- |

<sup>a</sup>Table provides mean (standard deviation; range); 7 participants were not included in the size analyses due to missing MI and 2 participants' MI could not be measured due to poor quality of MRI scan; Mann-Whitney U Tests were used for significance testing due to the negative skew of the data and rank biserial correlation ( $r_{rb}$ ) was used to estimate effect size

**Table 3.** Step-wise linear regression model analysis of massa intermedia size (*n* = 408)

| | $\beta$ (se) | Bootstrapped 95% CI<sup>†</sup> | $t$ | $p$ | $R^2_{\text{adj}}$ |
| --- | --- | --- | --- | --- | --- |
| **Axial** |  |  |  |  |  |
| *Step 1* |  |  |  |  | .181 |
| Sex (Male)<sup>a</sup> | -0.62 (0.34) | [-1.30, 0.06] | -1.81 | .071 | -- |
| Age | -0.12 (0.01) | [-0.15, -0.08] | -8.14 | <.001 | -- |
| Diagnosis (SZ)<sup>b</sup> | -1.55 (0.31) | [-2.16, -0.93] | -4.94 | <.001 | -- |
| *Step 2* |  |  |  |  | .200* |
| Sex<sup>a</sup> | -0.68 (0.33) | [-1.34, -0.00] | -2.03 | .043 | -- |
| Age | -0.06 (0.02) | [-0.11, -0.01] | -3.04 | .002 | -- |
| Diagnosis (SZ)<sup>b</sup> | 1.12 (0.87) | [-0.73, 2.95] | 1.28 | .200 | -- |
| Diagnosis (SZ) x Age<sup>c</sup> | -0.09 (0.02) | [-0.16, -0.02] | -3.27 | .001 | -- |
| **Coronal** |  |  |  |  |  |
| *Step 1* |  |  |  |  | .121 |
| Sex (Male)<sup>a</sup> | -0.78 (0.25) | [-1.28, -0.29] | -3.13 | .001 | -- |
| Age | -0.07 (0.01) | [-0.09, -0.04] | -6.47 | <.001 | -- |
| Diagnosis (SZ)<sup>b</sup> | -0.8 (0.23) | [-1.26, -0.34] | -3.47 | <.001 | -- |
| *Step 2* |  |  |  |  | .125<sup>d</sup> |
| Sex (Male)<sup>a</sup> | -0.81 (0.25) | [-1.29, -0.31] | -3.23 | .001 | -- |
| Age | -0.05 (0.01) | [-0.08, -0.01] | -3.09 | .002 | -- |
| Diagnosis (SZ)<sup>b</sup> | 0.19 (0.65) | [-1.10, 1.55] | 0.30 | .765 | -- |
| Diagnosis (SZ) x Age<sup>c</sup> | -0.03 (0.02) | [-0.08, 0.01] | -1.64 | .101 | -- |

<sup>†</sup> 5,000 bootstrapping model was used to generate 95% confidence interval
<sup>a</sup> Reference Category is Female
<sup>b</sup> Reference Category is Healthy Individual
<sup>c</sup> Reference Category is Healthy Individual x Age
\* The addition of a Diagnosis x Age interaction (Step 2) provides a significantly better fit than Step 1 ( $\Delta R^2_{\text{adj}} = .019$ , $p = .001$ )
<sup>d</sup> The addition of a Diagnosis x Age interaction (Step 2) did not provide a significantly better fit than Step 1 ( $\Delta R^2_{\text{adj}} = .004$ , $p = .1$ )

After adding the Diagnosis x Age interaction term in Step 2, the effect of diagnosis was no longer significant in both planes (Table 3). As shown in Figure 3, this was likely due to a more pronounced negative effect of age on MI lengths in SSD, though, the interaction term was significant only in the axial plane (axial: *p* = .001, *d* = -0.33; coronal: *p* = .101, *d* = -0.16). Notably, the significant and negative effect of age persisted in Step 2 in both planes (axial: *p* = .002, *d* = -0.30; coronal: *p* = .002, *d* = -0.31) and sex remained a significant effect in the coronal plane, with male sex associated with smaller MI length (*p* = .001, *d* = -0.32).

**Figure 1.**
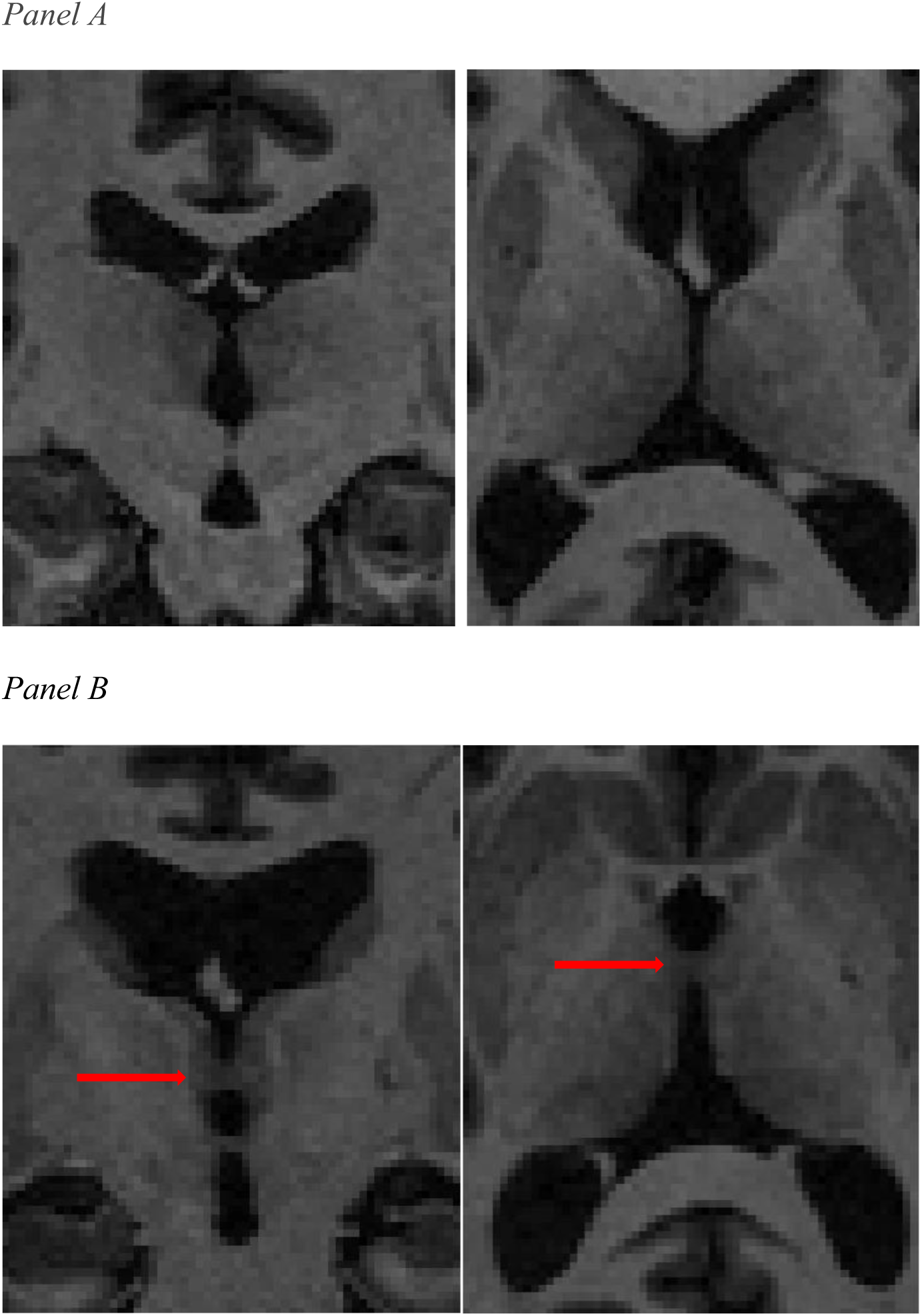
*Panel A:* Coronal (left) and axial (right) planes of an individual with missing MI. Only cerebrospinal fluid can be seen between the left and right thalamus. *Panel B:* Coronal (left) and axial (right) views of an individual with a MI (indicated by red arrows).

**Figure 2.**
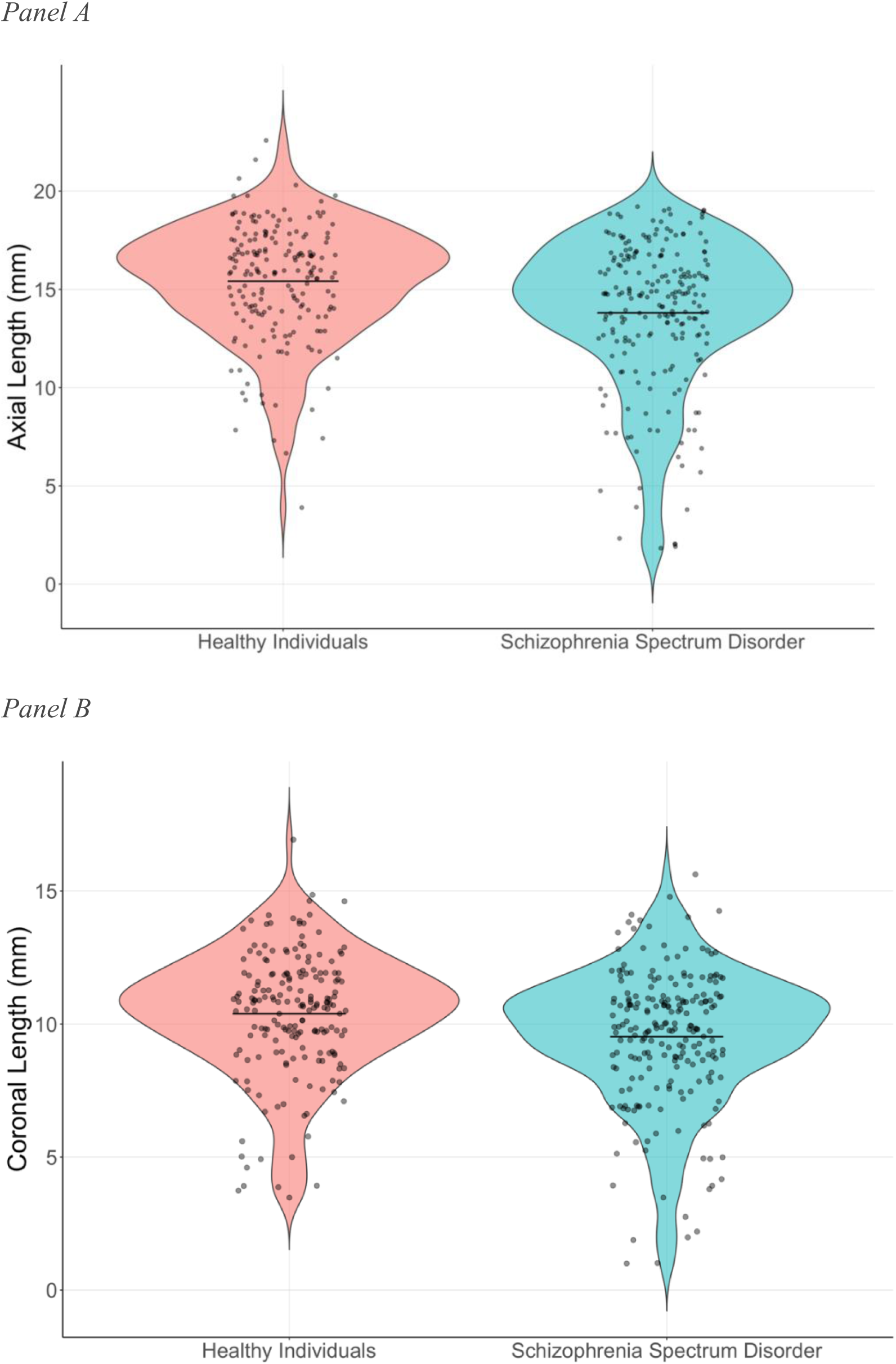
Violin plots of axial (Panel A) and coronal (Panel B) lengths in the schizophrenia spectrum disorder and healthy individual groups (*n* = 408). Bolded lines represent the mean MI size in each group.

**Figure 3.**
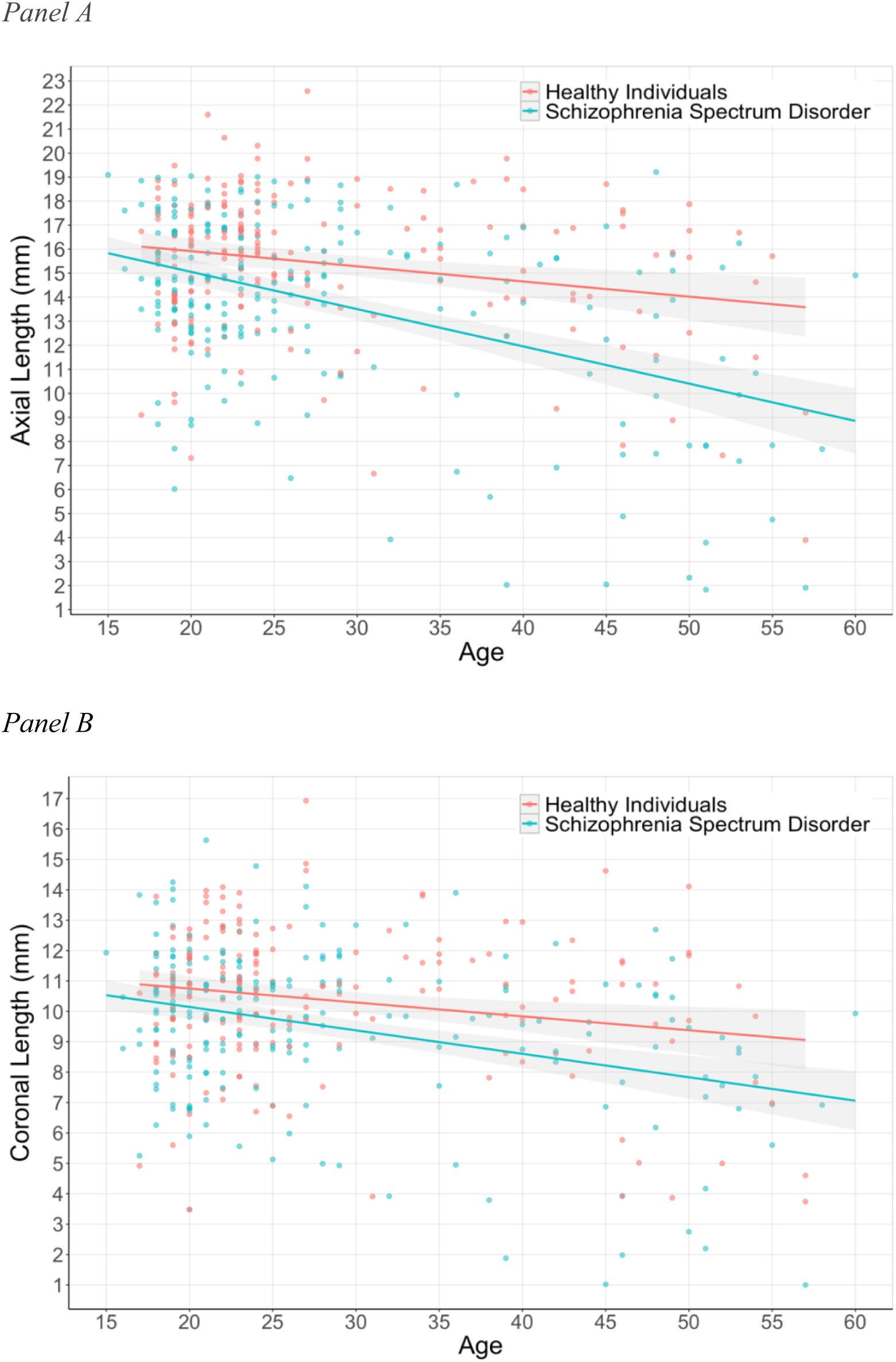
The association between age and axial MI length (Panel A) is significantly more negative in the schizophrenia spectrum disorder group than the healthy individual group (*n* = 408). The shaded area represents standard error.

### 3.2 Massa intermedia size and neurocognition

MI size (axial and coronal) was not a significant predictor of SCIP global cognition in Step 1 (Global Cognition ∼ MI Size + Age + Diagnosis). The addition of a Diagnosis x MI Size interaction in Step 2 (Global Cognition ∼ MI Size + Age + Diagnosis + Diagnosis x MI Size) did not significantly improve model fit. The SCIP subscales were not investigated in separate models because of the lack of association between MI Size and global cognition when controlling for the effects of age and diagnosis.

### 3.3 Massa intermedia morphology and incomplete hippocampal inversion

Chi-square test of independence found that IHI was significantly more common in individuals with missing MI (*χ*^2^=9.18, *p* = .002, Cramer’s V = .149). 5 of the 7 individuals with missing MI also had IHI. IHI presence was then added as a main effect to the linear regressions described above to investigate its effect on MI size (Step 3: MI size ∼ Sex + Age + Diagnosis + Diagnosis x Age + IHI). Including IHI presence did not significantly improve model fit in the axial plane. However, IHI presence did improve model fit in the coronal plane (ΔR^2^ = .008, *p* = .029). Pairwise comparisons revealed that those with IHI (Figure 4) had significantly shorter MI length in the coronal plane (β=-0.60, *t* = -2.19, SE = 0.27, *p* = .029, CI [-1.14, -0.07], *d* = -0.21). Spearman correlations were also run to examine continuous relationships between IHI scores (0-10) and MI size. No significant correlations between these variables were identified.

**Figure 4.**
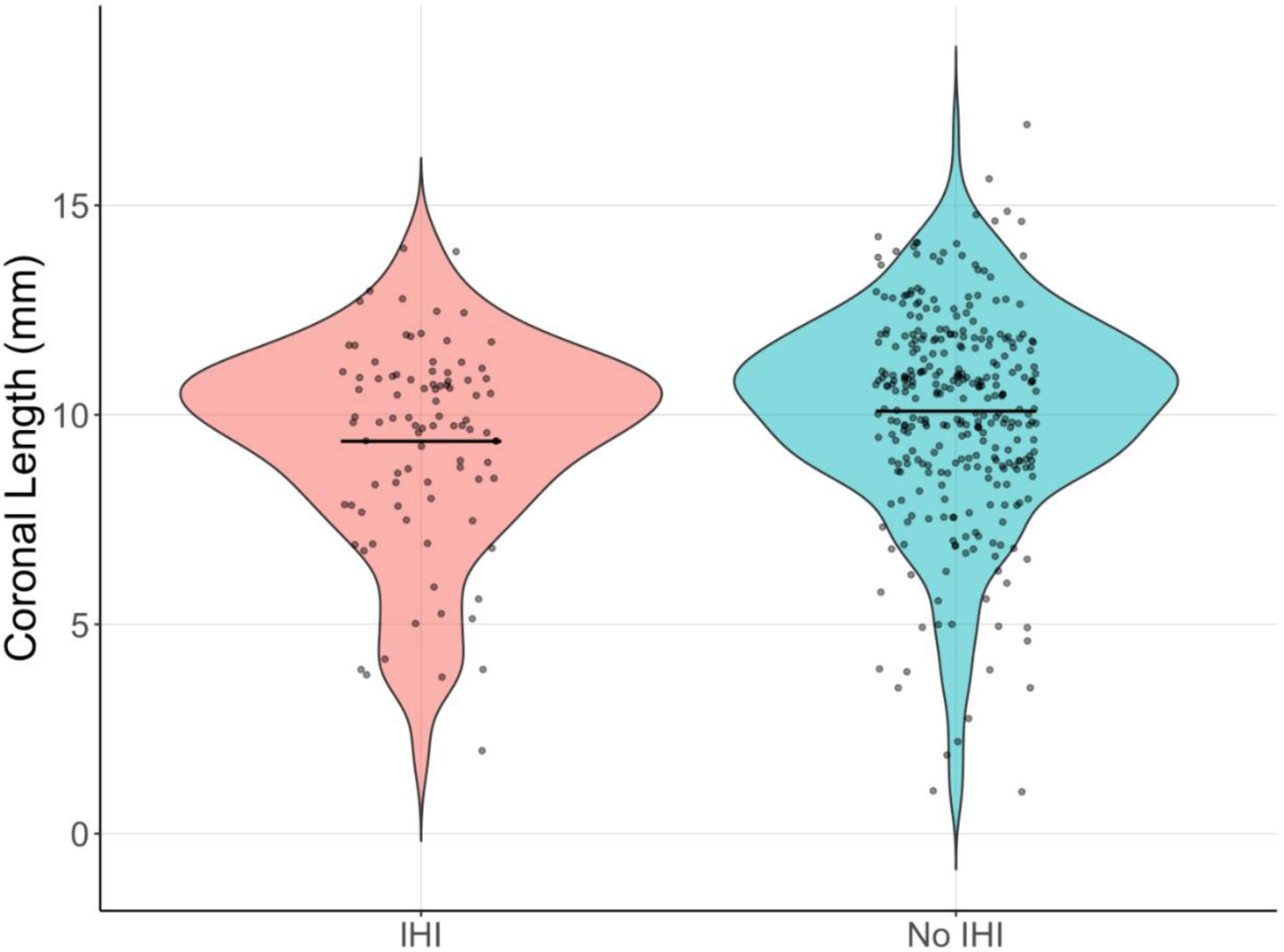
Coronal length among participants with and without incomplete hippocampal inversion (*n* = 408). Bolded lines represent mean coronal length in each group.

## 4. Discussion

Our study of 417 participants shows that missing MI is not more prevalent in individuals with schizophrenia spectrum disorders. The size of the MI was shown to be smaller in SSD than in healthy individuals. After accounting for a Diagnosis x Age interaction, we found that this smaller size may be caused by a more pronounced decline in size with age in SSD. This raises concerns about the reliability and validity of MI size measured in adulthood as an indicator of atypical neurodevelopment in SSD. Rather, smaller MI size could be a *neuroprogressive* feature of the disorder. Despite these observations regarding MI size in SSD, no significant relationships between MI morphology and neuropsychological functioning were identified in our models. Finally, although missing MI was not more common in SSD, it was more likely to occur in individuals with IHI. Moreover, the presence of IHI was associated with smaller MI in the coronal plane. This association between two markers of atypical brain development suggest a shared neurodevelopmental disruption in the second trimester.

Missing MI was not more prevalent in SSD. Since the MI forms in the second trimester (Rosales et al., 1968) our finding is not consistent with the neurodevelopmental hypothesis of schizophrenia (Marenco and Weinberger, 2000; Weinberger, 1987). This finding is in contrast to a meta-analysis conducted by Trzesniak et al. (2011), which found missing MI to be twice as likely in schizophrenia.

This discrepancy is likely partially attributable to the heterogeneity of methods used by the studies included in Trzesniak et al.’s meta-analysis. For instance, many had slice thickness’ greater than 1mm (Agarwal et al., 2008; Erbaǧcı et al., 2002; Snyder et al., 1998) and magnetic field strengths less than 3T (Ceyhan et al., 2008; Nopoulos et al., 2001; Takahashi, Suzuki, et al., 2008). Others marked the MI present only if it could be identified in at least 3 contiguous slices (Takahashi and Suzuki, et al., 2008; Takahashi and Yücel, et al., 2008). If these same methods were used in our study, it is likely that the rate of MI missingness in our sample would increase significantly, as we had many participants with small MIs.

It is also worth noting that it has been common to find no difference in missing MI between schizophrenia and healthy individuals. 6 of the 11 studies in Trzesniak’s et al.’s meta-analysis and 2 of the 3 most recent studies of the MI in schizophrenia (Landin-Romero et al., 2016; Takahashi et al., 2017; Trzesniak et al., 2012) did not find an increased prevalence of missing MI. These findings, in addition to the null result found in our large sample of 417 participants, suggests that missing MI is not a common neurodevelopmental feature of SSD.

Smaller MI appears to be a consistent morphological feature of SSD. This size difference occurred in both planes in our study (Table 2) and has also been found in other investigations of the MI (Takahashi et al., 2017; Trzesniak et al., 2012; Trzesniak, et al., 2011). Our results, however, indicate that smaller MI in SSD may not be a marker of atypical development. A large community sample (*N* = 1,410) of participants aged 2 months to 93 years indicated that the MI is not static in size across the lifespan, with the structure decreasing in size with age up to 70 (Borghei and Piracha, et al., 2021). A similar negative relationship with age was also found across our sample, with the decrease in size with age being more pronounced in SSD (Figure 3). In fact, when the Diagnosis x Age interaction was included in Step 2 of our linear model (Table 3), the main effect of diagnosis was no longer significant. This indicates that the diagnosis effect found in our study and others could be due to a neuroprogressive process (or perhaps accelerated aging), rather than a neurodevelopmental effect. Longitudinal research is needed to confirm this finding. If confirmed, neuroprogressive decline in MI size would be added to a growing list of neuroanatomical features that deteriorate faster in the lifetimes of individuals with schizophrenia (Cropley et al., 2017; Jiang et al., 2018; Lewandowski et al., 2020; Stone et al., 2022).

Modest associations between MI morphology and neuropsychological functioning have been identified in humans (Borghei et al., 2020; Damle et al., 2017; Trzesniak et al., 2016; Vidal et al., 2024). Our study appears to be the first to examine relationships between MI morphology and cognition in SSD. While our findings did not identify significant associations between the two, further investigations examining their relationship could be justified given the subtle associations with cognition in other human samples. For instance, SSD samples with larger rates of absent MI could allow researchers to examine the differential impact MI presence may have on cognition in the disorder. A similar analysis conducted by Trzesniak et al. (2016) in mesial temporal lobe epilepsy found patients without a MI had poorer verbal memory, executive control/attention, and long-term memory scores.

IHI occurs between gestational weeks 20-30 (Raininko and Bajic, 2010), is more likely to occur in preterm neonates (Bajic et al., 2010), and is the result of an arrest in brain development (Bajic et al., 2012). Experiencing this neurodevelopmental disruption made it more likely to have another type of atypical brain development in our study, missing MI. An investigation of pediatric patients with and without midline brain abnormalities identified a similar finding, as children with IHI had a higher likelihood of having missing MI (Whitehead and Najim, 2020). This commonality between IHI and missing MI suggests a shared disruption in brain development in the second trimester. It is currently unclear what this shared disruption could be, especially given that the MI forms before (13-14 gestational weeks) the hippocampal inversion process. Furthermore, we also identified an effect of IHI presence in our linear model, which indicated that individuals with IHI had smaller coronal lengths. The causal connection between IHI and smaller MI size is also not clear, and requires additional research to elucidate how this malformation of the hippocampus impacts other midline structures.

Our study has three significant limitations. First, we identified only 7 cases of missing MI in a sample of 417 participants. This resulted in a smaller rate of missing MI in both SSD and healthy individuals than has been found in other studies. It is unlikely, however, that there is something unusual about our sample or methods. Our MI size measurements were similar to findings from a large sample of healthy individuals (Borghei et al., 2020), and we replicated findings from other studies that found negative associations between age, male sex, and MI size (Allen and Gorski, 1991; Borghei et al., 2021; Damle et al., 2017; Takahashi et al., 2017). Moreover, variability in neuroimaging parameters and methods has led to a wide range of missing MI rates (approximately 4-25%) reported in the general population (Borghei et al., 2021; Damle et al., 2017; Nopoulos et al., 2001; Takahashi et al., 2017).

Second, our investigation of the MI was cross-sectional. Although reductions in MI size with age have been found in other large cross-sectional studies (Borghei, et al., 2021), it is still not certain that this structure undergoes an involution throughout the lifespan. Longitudinal research of the MI is required to validate the involution of this structure with age and an accelerated decline in its size in schizophrenia. Third, we did not collect information about obstetric complications in our participants. Obstetric complications increase schizophrenia susceptibility (Cannon et al., 2002) and significantly affect thalamic development (Ball et al., 2012; Volpe, 2009). Future studies should collect obstetric information to explore the relationships between MI, IHI, and complications in utero.

In conclusion, our large study of 417 participants did not find evidence that supports MI morphology (missingness, size) as a neurodevelopmental feature of SSD. On the contrary, we found evidence to support MI size as a *neuroprogressive* feature of the disorder, with its axial length potentially decreasing at an accelerated rate compared to healthy individuals. Longitudinal research is needed to confirm this finding. Although we did not find an increased prevalence of missing MI in SSD, this atypical midline feature was more common in individuals with IHI. Additional research is needed to investigate the possibility of a shared neurodevelopmental disruption in the second trimester that is responsible for these two features of atypical brain development.

